# Flexible parsing, interpretation, and editing of technical sequences with splitcode

**DOI:** 10.1101/2023.03.20.533521

**Authors:** Delaney K. Sullivan, Lior Pachter

**Affiliations:** UCLA-Caltech Medical Scientist Training Program, David Geffen School of Medicine, University of California, Los Angeles, Los Angeles, CA, 90095, USA; Division of Biology and Biological Engineering, California Institute of Technology, Pasadena, CA, 91125, USA; Department of Computing and Mathematical Sciences, California Institute of Technology, Pasadena, CA, 91125, USA

## Abstract

Next-generation sequencing libraries are constructed with numerous synthetic constructs such as sequencing adapters, barcodes, and unique molecular identifiers. Such sequences can be essential for interpreting results of sequencing assays, and when they contain information pertinent to an experiment, they must be processed and analyzed. We present a tool called splitcode, that enables flexible and efficient parsing, interpreting, and editing of sequencing reads. This versatile tool facilitates simple, reproducible preprocessing of reads from libraries constructed for a large array of single-cell and bulk sequencing assays.

**Availability and Implementation:** The splitcode program is free, open source, and available for download at http://github.com/pachterlab/splitcode.

## Introduction

The reads that result from next-generation sequencing libraries can contain many types of synthetic constructs, or technical sequences, including adapters, primers, indices, barcodes, and unique molecular identifiers (UMIs) (Kebschull and Zador 2018; Kivioja et al. 2011; Martin 2011; Melsted, Ntranos, and Pachter 2019; Johnson, Venkataram, and Kryazhimskiy 2023; Booeshaghi, Chen, and Pachter 2023). These oligonucleotide sequences are defined by the technicalities of sequencing based assays and experiments, with each sequence being either a completely unknown sequence, a known sequence, or an unknown sequence that is a member of a set of known sequences. There are many read preprocessing tools for editing and extracting information from such sequences, including the widely-used tools cutadapt (Martin 2011), fastp (Chen et al. 2018), and Trimmomatic (Bolger, Lohse, and Usadel 2014) for adapter and quality trimming, UMI-tools (Smith, Heger, and Sudbery 2017) and zUMIs (Parekh et al. 2018) for UMI processing, BBTools (Bushnell 2014; Bushnell, Rood, and Singer 2017) and reaper (Davis et al. 2013) for more general filtering operations, INTERSTELLAR for read structure interpretation (Kijima et al. 2023), among many other tools (Roehr, Dieterich, and Reinert 2017; Kong 2011; Battenberg et al. 2022; Liu 2019). However, no one tool can adequately address all technical sequence preprocessing tasks. Some methods, such as adapter trimming methods, can only remove identified technical sequences from reads but lack the ability to store information about technical sequences that are relevant to the provenance of the read. Other methods can extract and store technical sequences from reads but are limited to only extracting sequences at defined positions of defined lengths within reads, and may present limited options for handling variable position and variable length segments. Still other methods are designed for only a specific type of assay, such as single-cell RNA-seq. Technologies such as (long-read) SPLiT-seq (Rosenberg et al. 2018; Rebboah et al. 2021), SPRITE (Quinodoz et al. 2018, 2022), and Smart-seq3 (Hagemann-Jensen et al. 2020) contain complex, multifaceted technical sequences that currently are processed by custom scripts or specific use-case modifications to existing tools.

Here, we present splitcode which introduces versatile new features for general preprocessing needs. Splitcode is a flexible solution with a low memory and computational footprint that can reliably, efficiently, and error-tolerantly preprocess technical sequences based on a user-supplied structure of how those sequences are organized within reads. For example, splitcode can simultaneously trim technical sequences, parse combinatorial barcodes that are variable in length and inconsistent in location within a read, and extract UMIs that are defined in location with respect to other technical sequences rather than at a set position within a read. Moreover, splitcode can seamlessly interface with other command-line tools, including other read sequencing read preprocessors as well as read mappers, by streaming the pre-processed reads into those tools. Thus, splitcode can eliminate the need to write an entirely new file to disk at every step of preprocessing, a practice that currently results in inefficient use of time and disk space. Furthermore, splitcode can stream reads into itself, enabling multiple preprocessing steps to be performed in sequence for more complicated assays.

## Results

### Framework and Usage

We refer to the synthetic constructs, or technical sequences that can be identified in reads as tags. Tags are described in the splitcode config file with several parameters including a tag ID, the sequence itself, the error-tolerance for identifying that tag, and options such as where the tag might be found within sequencing reads and conditions under which the tag should be searched for. A collection of tags forms a barcode, which can be used to demultiplex reads according to the tags identified within a read. Within the config file, a user can also specify extraction options to delineate how certain subsequences within reads should be extracted. Subsequences can be extracted by using tags as anchor points or can be extracted at user-defined positions within reads. This feature is particularly useful for unique molecular identifier (UMI) sequences which are generally unknown sequences that exist at defined locations within reads. Additionally, in the config file, a user can specify read editing options including trimming and whether identified tags should be replaced with a particular sequence. Thus, identified technical sequences can be modified or trimmed *in situ*. Taken together, these array of options make it possible for splitcode to parse data from a large variety of sequencing assays, including those with many levels of multiplexing (**Figure 1**).

**Figure 1:**
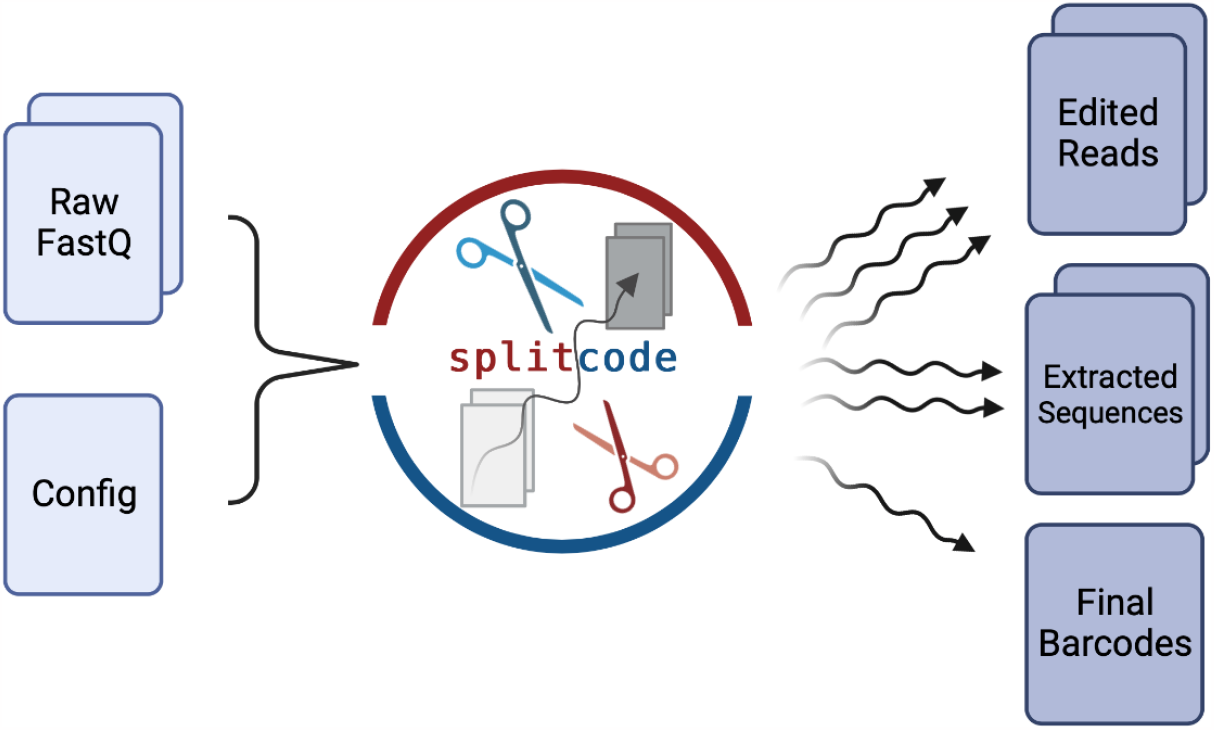
Overview of the splitcode workflow. The splitcode program takes in a set of FASTQ files and a user-specified config file, which serves as a recipe describing how the reads should be parsed. The user executes splitcode on the command-line, specifying command-line options on how the output should be formatted. The output consists of one or more of the following: the original FASTQ files (possibly edited), the extracted sequences (e.g. UMI sequences which are unknown and need to be extracted by using location information or anchor points), and the final barcodes which are unique for each combination of identified tags. The output may take the form of FASTQ files, gzip-compressed FASTQ files, or interleaved sequences directed to standard output, depending on what the user specifies.

Following construction of the config file (**Figure 2**), users can supply the config file to the splitcode program on the command-line. Users can further specify the output options for how the final barcode, the (possibly edited) reads, the extracted subsequences should be outputted. The program presents many options for outputting reads, allowing seamless integration with many downstream tools. Importantly, the output can be interleaved and directed to standard output, which can then be directly piped into tools (including splitcode itself if another round of read processing is needed) that support such input. This feature makes it possible to send processed reads directly to a read mapper, therefore eschewing the inefficiencies of creating large intermediate files on disk.

**Figure 2:**
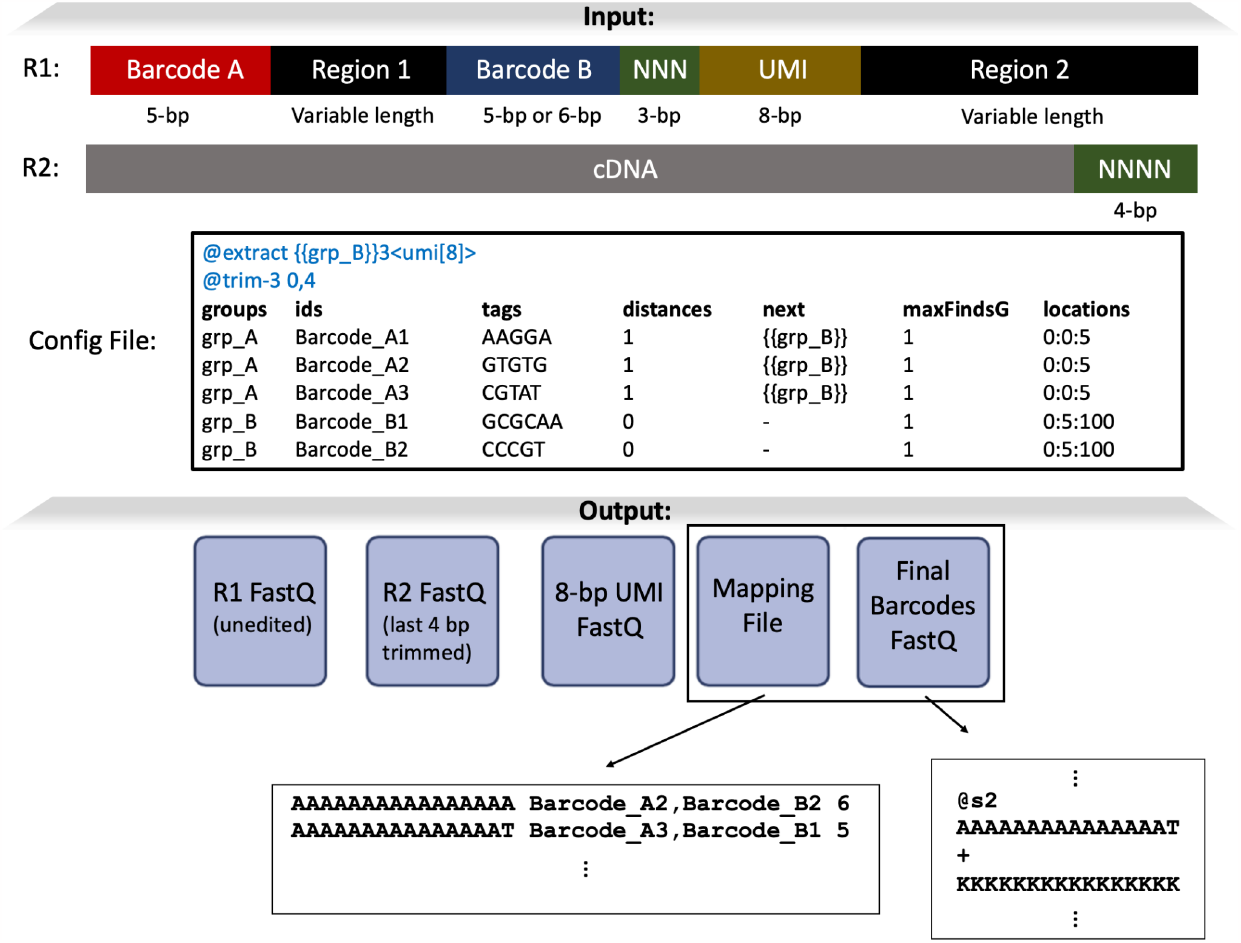
Example of splitcode usage. The structure of the reads from this hypothetical sequencing technology contains multiple regions that need to be parsed, including some of variable length. In the config file, each region that needs to be parsed is organized into “groups” and each group contains multiple tags. The tags in the grp_A group have the value 1 in the “distances” column, meaning a hamming distance 1 error tolerance. The values in the “next” column indicate that after a grp_A tag (i.e. Barcode_A1, Barcode_A2, or Barcode_A3) is found, we should next search only for tags in the grp_B group. The “maxFindsG” values of 1 mean that the maximum number of times a specific group can be found is 1 (e.g. after finding a tag in grp_A, stop searching for tags in grp_A). The “locations” for grp_A tags have the value 0:0:5, meaning that the tag is found in file #0 (i.e. the R1 file) within positions 0-5 of the read; for grp_B tags, splitcode searches file #0 within positions 5-100. In the header of the config file, the @extract option contains an expression indicating that we should extract an 8-bp sequence, which we name umi, 3 bases following identification of a grp_B tag. The supplied @trim-3 option means that only 3′-end trimming of 0 bases and 4 bases of the R1 file and the R2 file, respectively, should be performed. As output, the “Final Barcodes” FASTQ file contains a sequence uniquely identifying a combination of tags and the mapping file allows us to map the final barcode sequence back to the tag combination (the numbers in the right-most column of the mapping file represent how many reads that tag combination was found in). Finally, it is important to note that this is simply one of many ways to parse this read structure with splitcode and users can configure the options how they save fit. Further, users can also customize the output options (for example, users can choose to output reads that contain both grp_A and grp_B tags into one set of files and direct all other reads into a separate set of files, and users can choose whether to output the 8-bp UMI sequence into an independent file or to put it in the FASTQ header of the outputted reads as a SAM tag).

### Capabilities

The splitcode program has many options, some of which can be supplied in the config file and others of which (namely the output options) must be supplied on the command line. In the config file, a user can specify “sequence identification” options for finding tags in reads as well as editing reads *in situ* based on identified tags as well as “read modification and extraction” options for general read trimming and extracting UMI-like sequences. The latter option group is supplied in the header of the config file while the “sequence identification” options are supplied as tab-separated values in a tabular format in the file, an example of which is shown in Figure 2. A list of some of the splitcode config file options is exhibited in Table 1.

**Table 1:**
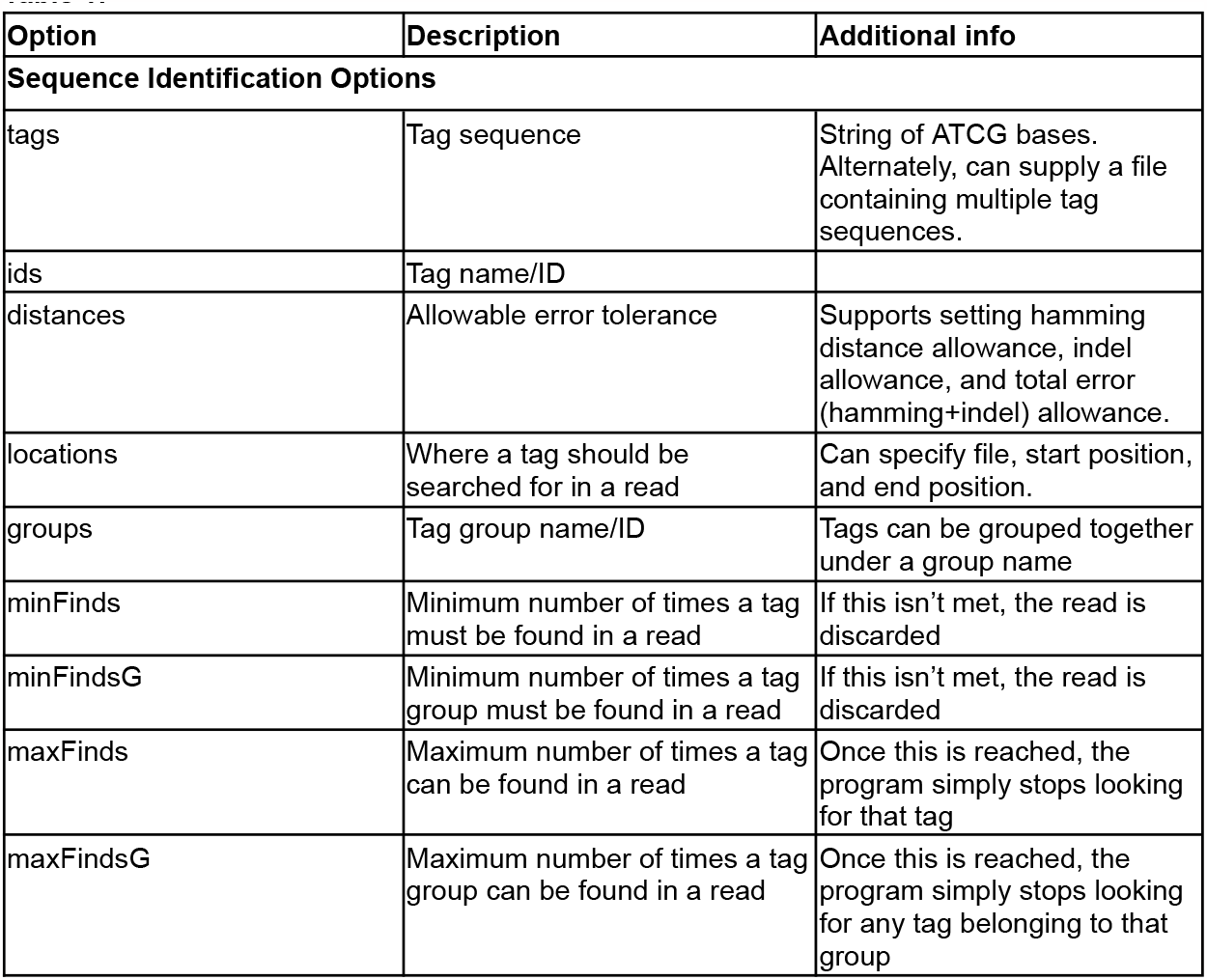
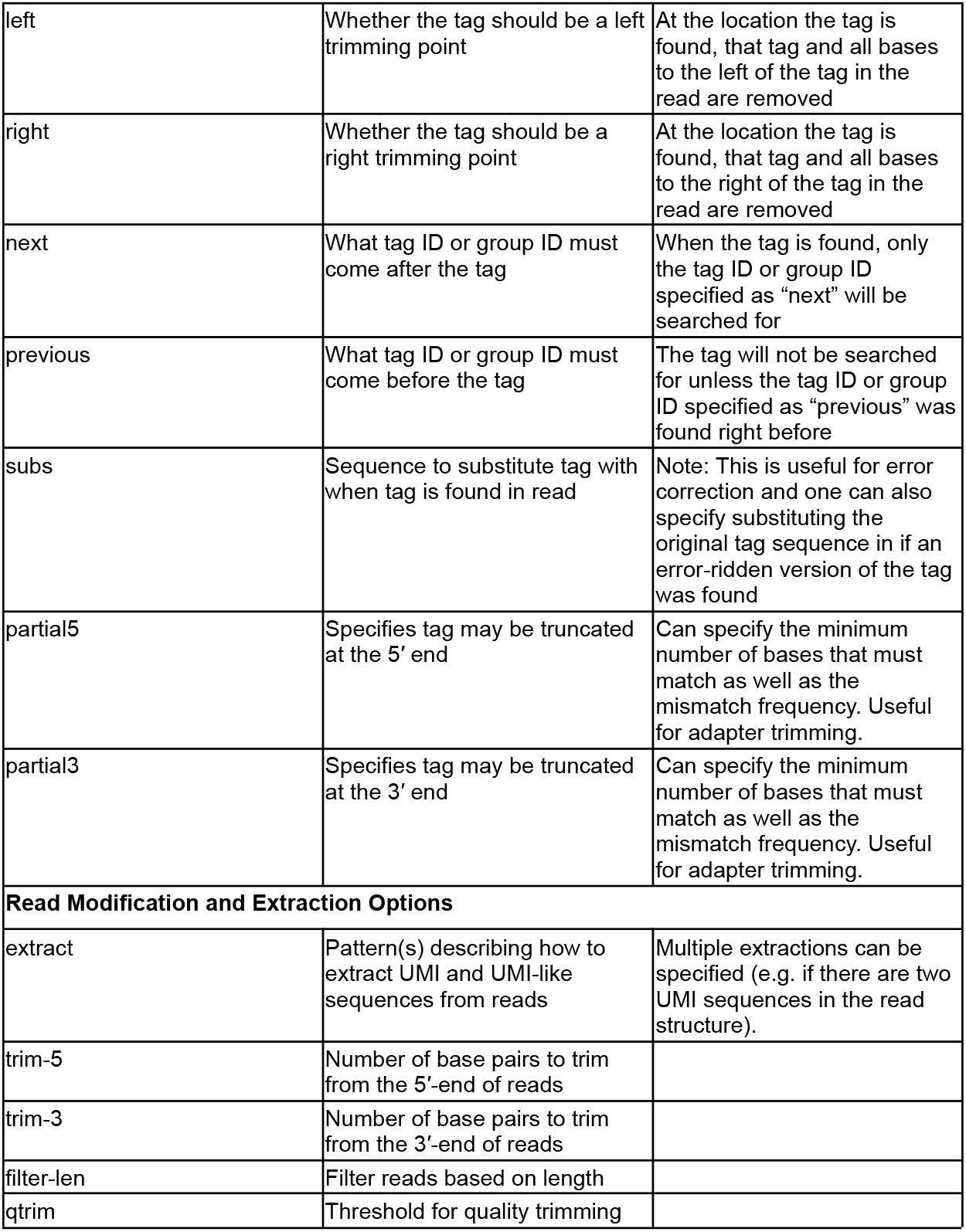

The features specified in **Table 1** relate to read editing and tag identification. Downstream of the tag identification process, there are more options to further process identified tags. For example, using the --keep and --keep-grp command-line options, a user can specify combination(s) of tags or tag groups that should be retained and only those reads will be kept. Likewise, using the --discard and --discard-grp command-line options, a user can specify a combination(s) that should be discarded and those reads will be discarded. Furthermore, using the --keep and --keep-grp options, a user can specify specific combinations to be outputted into specific files, enabling demultiplexing based on barcodes or barcode combinations.

A graphical user interface (GUI) for splitcode facilitates the usage of splitcode (Figure 3). This GUI exists as a web page and helps a user create a config file which can then be downloaded. Additionally, this GUI enables live testing of configuration options on user-supplied sample sequences.

**Figure 3:**
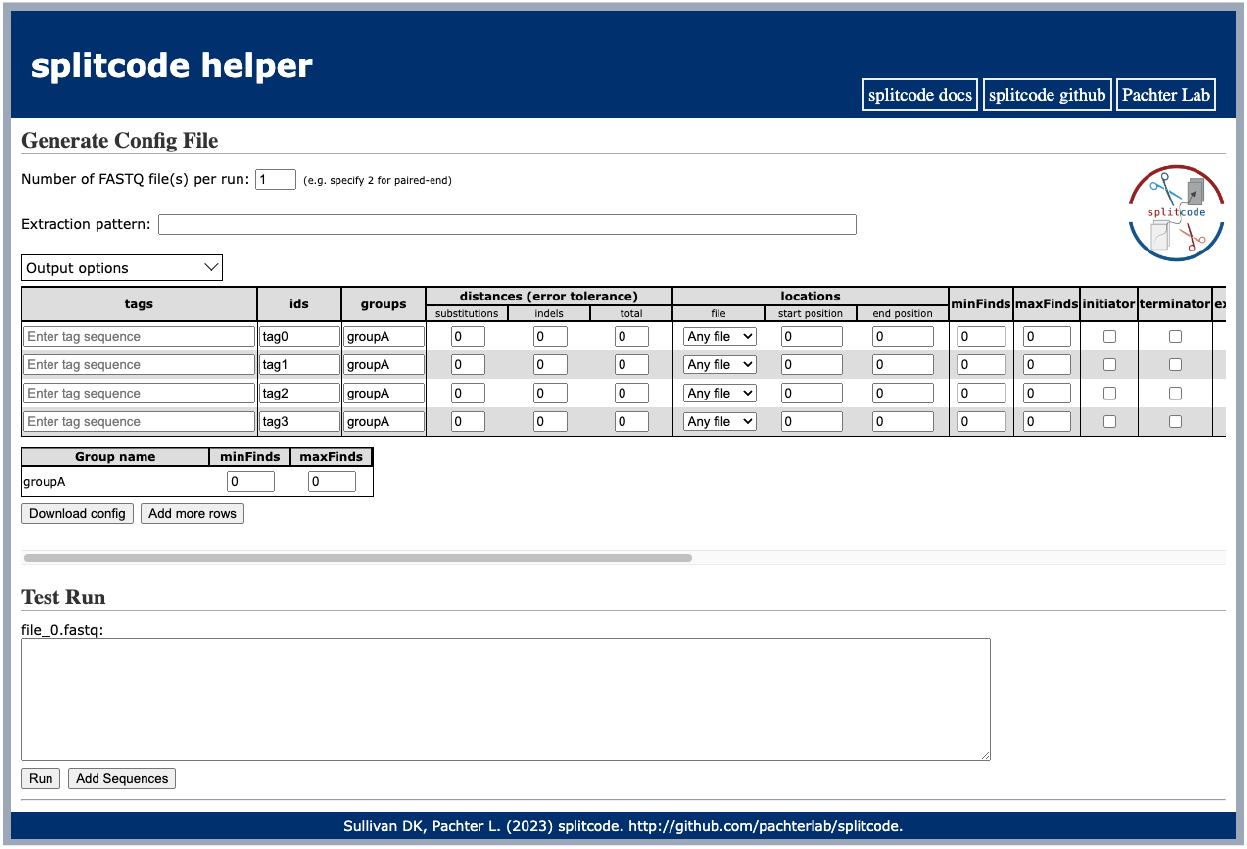
The splitcode graphical user interface (GUI). The GUI can be viewed in a web browser and is designed to facilitate creation of the splitcode config file and navigation of output options. The GUI also features live testing of the splitcode program on user-supplied sample sequences in FASTQ format.

## Discussion

The preprocessing of FASTQ files is an important first step in bioinformatics pipelines. This step is frequently inefficient, involving multiple steps with the creation of large intermediate files or writing and running of custom unoptimized scripts which can be challenging with large-scale sequencing data. Splitcode alleviates some of these inefficiencies via a modular and flexible design to effectively and efficiently handle intricate, hierarchical read structures produced by technologies with many layers of multiplexing. While many of splitcode’s features overlap with those of existing bioinformatics software, splitcode is not intended to fully recapitulate all the features of existing tools or to replace or outperform any one tool. Rather, splitcode is intended to serve as one additional, flexible and versatile tool in a bioinformatics arsenal, and has been designed to be interoperable with other tools. splitcode operates not as an alignment algorithm, but on a principle of dictionary lookups. In this approach, technical sequences along with their permissible mismatches are cataloged in a hash table. This makes splitcode apt for scenarios requiring identification, interpretation, and modification of short sequences within reads, and it effectively manages extensive lists of lookup sequences. Algorithms like cutadapt which use dynamic programming score matrix to optimize alignment, are more suitable for cases, such as general adapter trimming, that require finding the best possible alignment between two sequences or for finding long technical sequences (in which case, storing the allowable mismatches in a hash table is computationally infeasible). We anticipate that splitcode will be used in tandem with other preprocessing tools to provide an effective solution for many bioinformatics needs. Furthermore, we expect that splitcode will continue to expand in functionality based on user feedback, user needs, and possibly the introduction of more complicated read structures that may arise from the development of novel sequence census assays.

## Methods

### Tag Sequence Identification

Each sequence in the config file along with all sequences within the sequence’s allowable hamming distance and/or indel error tolerance is indexed in a hash map. Each sequence is associated with the tag(s) from which it originated. Reads in FASTQ files are scanned from start to end to identify tags based on hash map lookups. Additionally, users can specify locations and conditions within which a specific tag may appear and only tags satisfying such conditions are identified. Further, by restricting tag identification to only specific regions of reads, the number of hash map queries is reduced therefore improving runtime.

### Final Barcode Sequences

Each combination of tags is assigned a numerical ID, which begins at 0 and is incremented for every newly encountered combination. Each numerical ID, a 32-bit unsigned integer, can be converted to a unique 16-bp final barcode sequence by mapping each nucleotide to a 2-bit binary representation as follows: A = 00, C = 01, G = 10, T = 11. It follows that the numerical ID can be represented in nucleotide-space based on the integer’s binary representation. For example, the numerical ID 0 is AAAAAAAAAAAAAAAA, the numerical ID 1 is AAAAAAAAAAAAAAAT, and the numerical ID 30 is AAAAAAAAAAAAACTG. This interconversion between numerical IDs and nucleotide sequences facilitates simplifying complex barcodes.

### Software

The splitcode software is written in C++11 and is freely available and open source under the BSD-2 clause license. The framework for splitcode is a C++ header file making the direct incorporation of splitcode into a software project that involves processing sequencing reads possible. The GUI for the software is implemented as an HTML webpage and uses Emscripten for compilation of the software to WebAssembly. Documentation for the software is available at https://splitcode.readthedocs.io/.

## Contributions

D.K.S. conceived of the work, developed the methods and software, and drafted the manuscript.

L.P. supervised the work. Both authors reviewed and edited the manuscript.

## Acknowledgments

We thank Benjamin T. Yeh (Caltech) and the laboratory of Mitchell Guttman (Caltech) for discussions which motivated this project. Some of the splitcode source code is derived from source code written by Páll Melsted (University of Iceland), and we are grateful to him for sharing his source code with us. Thanks to A. Sina Booeshaghi for helpful discussions regarding *seqspec* and splitcode. Illustrations were created with BioRender: http://biorender.com.

## Funding

D.K.S. was funded by the UCLA-Caltech Medical Scientist Training Program (NIH NIGMS training grant T32 GM008042). L.P. was supported in part by the National Institutes of Health (NIH) grants U19MH114830 and 5UM1HG012077-02. The authors declare no conflicts of interest.

## Code Availability

The splitcode software is available at http://github.com/pachterlab/splitcode. The version of the splitcode software referred to throughout this paper is version 0.29.0.

## Notes

### Competing Interest Statement

The authors have declared no competing interest.

### Summary of Updates

Improved the text

